# An Electromechanical Relay-like Multi-enzymatic System Gates c-di-GMP-Dependent Cell Fate in a Cyanobacterium

**DOI:** 10.1101/2025.11.04.686522

**Authors:** Qing-Xue Sun, Min Huang, Xiaoli Zeng, Cheng-Cai Zhang

## Abstract

Cyclic-di-GMP is a ubiquitous bacterial second messenger that regulates diverse cellular processes. Although many bacteria harbor multiple enzymes for metabolism, the mechanisms underlying coordinated regulation remain unclear. Using the cyanobacterium *Anabaena* PCC 7120, we generate mutant strains with varying numbers of c-di-GMP metabolic genes deleted, including cdg^0^ (all degradation genes deleted) and cdg^max^ (all synthesis genes deleted). We found that cyclic-di-GMP is essential for both cell viability and size regulation. Quantitative analysis identified two critical physiological thresholds: one for maintaining normal cell size and a lower, lethal threshold required for survival. The 16 cyclic-di-GMP metabolic enzymes function as an electromechanical-like dual relay system, where different enzyme groups maintain cyclic-di-GMP homeostasis or activate SOS-like alarm responses when cyclic-di-GMP concentrations drop below the lethal threshold. These regulatory effects are mediated by the cyclic-di-GMP receptor CdgR, depending on the fraction of apo-CdgR form. This dual-threshold system enables dynamic cellular adaptation while preventing lethal consequences, representing a fundamental growth-survival trade-off strategy in living organisms.

## Introduction

cyclic di-GMP (c-di-GMP), a bacterial second messenger, governs a large spectrum of cellular processes (Enomoto *et al*, 2015; Enomoto *et al*, 2023; Hendrix *et al*, 2024; Jenal *et al*, 2017; Römling *et al*, 2013). The regulatory effects of c-di-GMP are mediated by binding to specific receptors, which exhibit remarkable structural and sequence diversity across bacterial species (Amikam & Galperin, 2006; Chou & Galperin, 2016; Hengge, 2009; Zeng *et al*, 2023). These regulatory effects can occur at multiple levels, including transcriptional, post-transcriptional, and post-translational regulation (Guo *et al*, 2025; Hengge, 2010; Jenal *et al*., 2017; Schäper *et al*, 2017; Wang *et al*, 2016).

The intracellular levels of c-di-GMP are finely regulated by two classes of enzymes: diguanylate cyclases (DGCs) and phosphodiesterases (PDEs) (Jenal *et al*., 2017). DGCs, characterized by a conserved GGDEF domain, are responsible for synthesizing c-di-GMP by the condensation of two GTP molecules (Jenal *et al*., 2017; Wassmann *et al*, 2007). Phosphodiesterases are responsible for the degradation of c-di-GMP. PDEs typically possess either an EAL domain or an HD-GYP domain, and hydrolyze c-di-GMP into linear pGpG or eventually into GMP (Christen *et al*, 2005; Jenal *et al*., 2017). Importantly, GGDEF, EAL, and HD-GYP domains are often found within the same protein, and their output activities can be regulated by various sensory domains that respond to a range of environmental signals (Agostoni *et al*, 2013; Enomoto *et al*., 2015; Enomoto *et al*., 2023; Yu *et al*, 2023). Additionally, DGCs often possess a c-di-GMP-binding inhibitory (I)-site involved in allosteric feedback inhibition (Yang *et al*, 2011). Consequently, the intracellular levels of c-di-GMP can be finely tuned in response to internal and external environmental cues.

Most bacterial genomes contain multiple genes related to c-di-GMP turnover (Agostoni *et al*., 2013). For example, *E. coli* and *Pseudomonas aeruginosa* contain 29 and 41 such genes, respectively (Agostoni *et al*., 2013). This genetic abundance implies sophisticated regulation mechanisms to control the c-di-GMP pools in response to various environmental cues. Some studies have revealed critical insights into c-di-GMP-mediated regulation (Abel *et al*, 2013; Chen *et al*, 2020; Isenberg *et al*, 2022; Kharadi *et al*, 2022; Solano *et al*, 2009). For example, in *Salmonella*, most GGDEF proteins are constitutively and independently expressed (Solano *et al*., 2009). In *Caulobacter crescentus*, c-di-GMP controls multiple developmental pathways, each with a distinct activation concentration (Abel *et al*., 2013). Additionally, in *Dickeya zeae* EC, 3 out of 19 c-di-GMP turnover proteins play dominant roles in modulating the global c-di-GMP pool (Chen *et al*., 2020). These findings collectively indicate the complexity and adaptability of c-di-GMP signaling networks in bacterial systems. However, it remains unclear why bacteria possess so many genes for c-di-GMP turnover and what the functional purpose and interplay of these genes are.

*Anabaena* sp. PCC 7120 (*Anabaena*) is a filamentous cyanobacterium that houses 16 genes for c-di-GMP metabolism (Agostoni *et al*., 2013) (Huang *et al*, 2021; Zeng & Zhang, 2022). We recently identified c-di-GMP as an intracellular proxy for cell size control (Sun *et al*, 2024) and discovered a highly conserved c-di-GMP receptor (CdgR), which modulates cell size through interaction with DevH *(Xu et al, 2025; Zeng et al., 2023)*. While individual deletion of the 16 genes led to the identification of CdgS as the primary DGC responsible for cell size regulation, by forming a two-component system with the histidine kinase CdgK, other single mutants had little noticeable phenotype (Huang *et al*., 2021; Neunuebel & Golden, 2008; Sun *et al*., 2024). Therefore, potential compensatory or redundant functions of the remaining 15 c-di-GMP metabolic proteins in cell size control and other physiological processes need to be elucidated.

To investigate how c-di-GMP turnover proteins modulate physiology in *Anabaena*, we methodically engineered two mutants: a c-di-GMP-deficient strain (*cdG^0^*) and a c-di-GMP-overproducing strain (*cdG^max^*) by sequentially deleting all 14 GGDEF-domain genes and all 8 EAL/HD-GYP domain genes, respectively. The deletion of such a large number of genes is unprecedented in cyanobacterial genetics and allowed us to determine the physiological functions of c-di-GMP. Several mutants that produce intermediate levels of c-di-GMP in this process were also obtained. By systematic analysis, we identified the dominant DGCs and PDEs in this cyanobacterium. We propose that the c-di-GMP metabolic enzymes function in a manner that simulates an electromechanical dual relay system to control the c-di-GMP hemostasis. In this system, thirteen enzymes constitute a baseline signal power, two acting as a responsive relay, and one as an emergency relay that is activated only when c-di-GMP concentration drops to a lethal level. These functional modules determine two concentration thresholds that dictate cell size and viability, respectively.

## Results

### Analysis of a mutant in which all 8 genes encoding c-di-GMP hydrolase were deleted demonstrates that high c-di-GMP levels do not affect cell growth and cell size

*Anabaena* possesses 16 proteins involved in c-di-GMP turnover (Agostoni *et al*., 2013; Sun *et al*., 2024). These include 8 proteins with GGDEF domains (DGC-only), 2 proteins containing EAL/HD-GYP domains (PDE-only), and 6 hybrid proteins that contain both types of protein domains (Dual-function) (Fig. S1A). Previous study revealed that neither deleting a single protein containing the EAL domain nor overexpressing the heterologous DGC YdeH had any observable impact on cell size or cell growth (Huang *et al*., 2021; Sun *et al*., 2024). This suggested the functional redundancy among the 8 c-di-GMP hydrolases (8 PDEs).

To overcome this redundancy and investigate the impact of high c-di-GMP levels in *Anabaena*, we systematically deleted all 8 PDE genes in frame based on CRISPR-Cpf1 (Niu *et al*, 2019), generating a PDE-null mutant (*8ΔPDE*, *cdG^max^*) and seven intermediate strains (*1ΔPDE*, *2ΔPDE*, *3ΔPDE*, *4ΔPDE*, *5ΔPDE*, *6ΔPDE*, *7ΔPDE* (Fig. S1B). Liquid chromatography-mass spectrometry (LC-MS) quantification of intracellular c-di-GMP revealed that a slight increase was observed upon deletion of the first single gene, *alr2306* (*1ΔPDE*), and the levels in the other 5 mutants (*2ΔPDE, 3ΔPDE, 4ΔPDE, 5ΔPDE, 6ΔPDE*) showed no significant difference from the WT strain (Fig. 1A). Deletion of the 7th PDE gene (*all4897*) triggered a sharp increase in c-di-GMP, and this level was further elevated in the *cdG^max^* strain, reaching a peak approximately 3.6-fold higher than WT following the deletion of *alr3170* (Fig. 1A). In contrast, *Δall3170* and *Δall4897* single mutants exhibited WT c-di-GMP levels (Fig. 1A), suggesting functional redundancy and cross regulation among these PDE enzymes. Next, we analyzed the cell morphology of all mutants and found they all displayed cell length and width similar to the WT strain (Fig.1B and Fig.S2). The *cdG^max^* strain also showed no growth defects under either BG11 or BG110 conditions, compared to the WT (Fig. S3). Together, these findings demonstrate that elevated c-di-GMP levels do not affect cell growth and cell size in *Anabaena*.

**Fig. 1.**
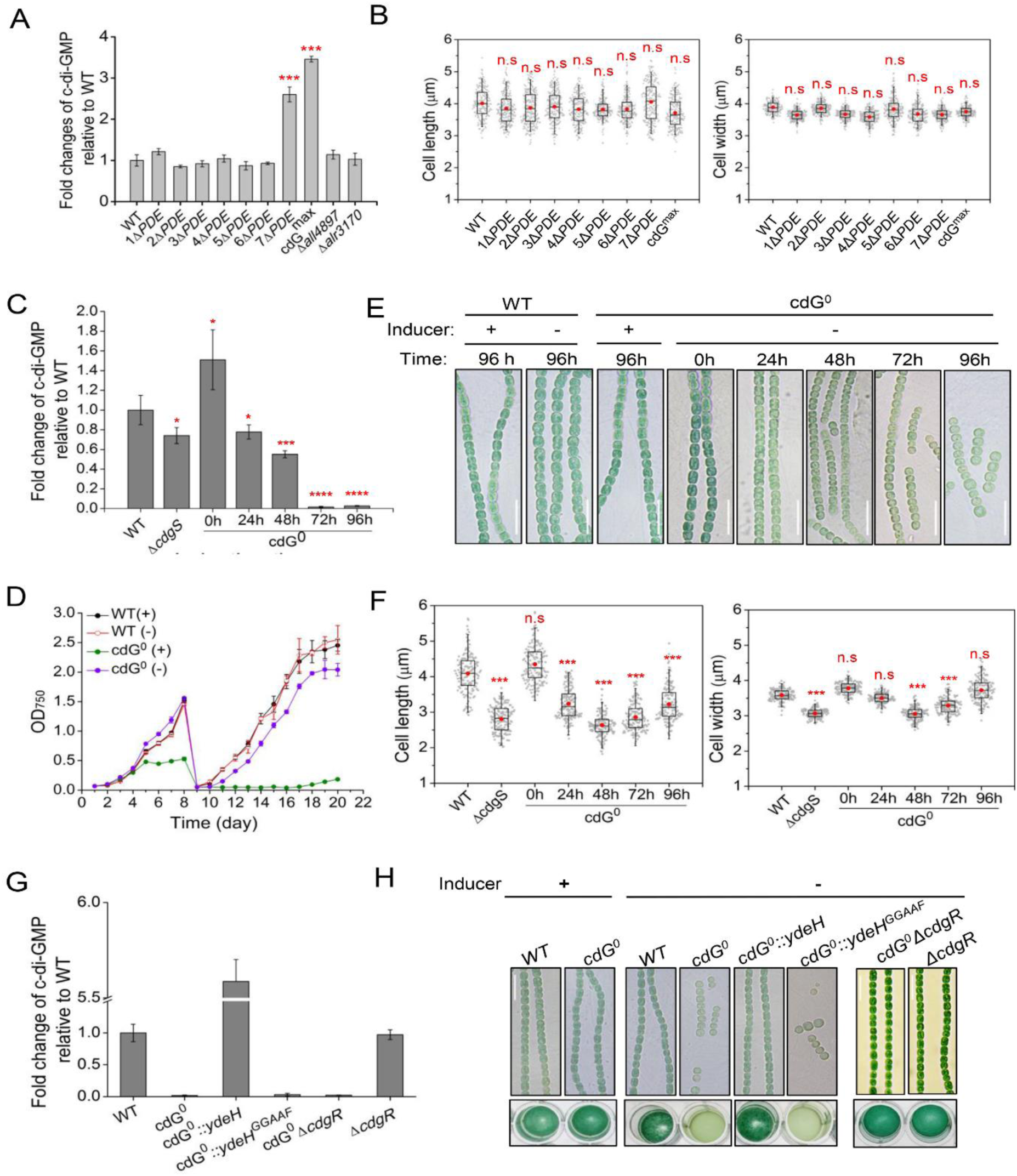
The phenotype and c-di-GMP level of all PDE deletion strains and the *cdG^0^* strain. (A) The intracellular c-di-GMP levels of the indicated strains. (B) Statistical analysis of cell length and cell width of the indicated strains based on images, as shown in Figure S2, using a box plot. (C) The intracellular c-di-GMP levels of the *cdG^0^* strain during the time course of Cu^2+^ and theophylline depletion. (D) The growth curves of the *cdG^0^* strain and WT strain in BG11 medium with (+) or without (-) Cu^2+^ and theophylline. Absorbance at 750 nm was measured at the indicated time points. To better remove traces of Cu^2+^ and theophylline, the cultures were re-inoculated for the second time in a medium free of Cu^2+^ and theophylline after 8 days of incubation. All values are shown as mean ± standard deviation, calculated from triplicate data. (E) Micrographs of *Anabaena* filaments of WT and *cdG^0^*strains cultured in BG11 medium with (+) or without (-) Cu^2+^ and theophylline at the indicated time points. Scale bars represent 15µm. (F) Statistical analysis of cell size parameters of the *cdG^0^* strain during the time course of Cu^2+^ and theophylline depletion. The cell length and cell width were measured based on images, as shown in (E). (G) The intracellular c-di-GMP level of the indicated strains at 96 h in BG11 medium without Cu^2+^ and theophylline. (H) Top panel: the micrographs of the indicated strains cultured in BG11 medium with (+) or without (-) Cu^2+^ and theophylline at 96 h. Bottom panel: The growth of different strains, as in the top panel tested in 24-well plates. All cultures started with a similar OD at 0.3 diluted from a pre-culture and imaged after 4 days of incubation. In (A), (C), and (G), the c-di-GMP concentrations of all mutants were normalized to the levels of the wild-type strain. The fold change numbers are shown as mean ± SD from three biological replicates. In (B) and (F),150-200 cells of each strain from three independent experiments were measured. The boxplots enclose the 25th and the 75th percentile, with the black line representing the median value, red dot representing the mean value. The statistical significance in comparison to WT was carried out by a two-sided Student’s t-test. The red asterisks indicate significance in comparison to WT as follows: *, p<0.05; **, p<0.01; ***, p<0.001. ****, p<0.000, n.s.: not significant (p >0.05). WT, wild type *Anabaena*.

### c-di-GMP is essential for cell survival

Previous studies demonstrated that the deletion of *cdgS* (strain *ΔcdgS* or 1*ΔDGC*) significantly reduced cell size (Neunuebel & Golden, 2008; Sun *et al*., 2024). Quantification analysis revealed that the intracellular c-di-GMP level was approximately 30 ± 11% lower than that in the WT strain (Fig. 1C). To explore the effects of additional c-di-GMP reduction and the roles of other DGCs in c-di-GMP homeostasis, we attempted to construct a DGC-null mutant (*cdG^0^*) by systematically deleting all 14 genes encoding GGDEF domain-containing proteins (Fig. S1C). While 13 of these genes were successfully deleted (13*ΔDGC*), the final gene, *all1219*, could not be inactivated in this background, while *all1219* alone could be inactivated as reported by two independent studies (Neunuebel & Golden, 2008; Sun *et al*., 2024). This suggests that c-di-GMP may be essential for the survival of *Anabaena*.

To overcome this challenge, we generated a conditional mutant by replacing the native promoter of *all1219* with the Cu^2+^ and theophylline (TP) inducible one (*CT* promoter) (Huang *et al*., 2021) in the *13ΔDGC* background strain, generating the *14ΔDGC* (*cdG^0^*) strain (Fig. S4A). When cultured in BG11 medium supplemented with 0.5 mM TP and 0.3 µM Cu^2+^, the *cdG^0^* strain exhibited cell growth (Fig. 1D) and cell size (Fig. 1E and 1F) comparable to those of the WT strain, with c-di-GMP levels approximately 1.4-fold higher than WT (Fig. 1C). Upon inducer removal, the intracellular c-di-GMP level gradually decrease, reaching an undetectable level by 72 h (Fig. 1C). Subsequently, by 120 h post-inductor removal, *cdG^0^* strain exhibited a complete growth arrest (Fig. 1C and Fig. S4B). These results demonstrate that the CT promoter could tightly regulate *all1219* expression, and confirm that c-di-GMP is essential for cell viability in *Anabaena*.

Consistent with previous reports (Sun *et al*., 2024), under conditions of c-di-GMP depletion (achieved by the removal of Cu²⁺ and TP), the average cell length and width of the *cdG^0^* strain gradually decreased over time (Fig. 1E and 1F). At 48 h after removing inducer, as the intracellular c-di-GMP dropped to 55 ± 11% of that found in the WT strain, the average cell length and width of the *cdG^0^* strain were reduced to 2.6 ± 0.32 µm and 3.0 ± 0.19 µm, respectively, compared to 4.1 ± 0.54 µm and 3.6 ±0.16 µm in WT strain (Fig. 1E and 1F). A slight recovery in the average cell length and width was observed at 72 h and 96 h (Fig. 1E and Fig. 1F), even though the c-di-GMP level had become undetectable (Fig. 1B). However, this recovery coincided with filament fragmentation and a gradual arrest in cell growth (Fig. 1E and 1F), suggesting the existence of a transient compensatory mechanism that ultimately fails to sustain normal cellular function.

To further confirm that the c-di-GMP reduction is responsible for the defects of the *cdG^0^* strain, we overexpressed a well-characterized heterologous DGC, YdeH, or its inactive variant YdeH^GGAAF^ (Huang *et al*., 2021; Zähringer *et al*, 2011) in this background, generating the strains *cdG^0^*::*ydeH* and *cdG^0^*::*ydeH^GGAAF^*, respectively. Upon the inducer removal, the expression of YdeH in the *cdG^0^*::*ydeH* strain resulted in the production of c-di-GMP levels approximately 5.6-fold higher than those in the WT strain, and it fully restored the cell size and cell growth defect of the *cdG^0^* strain (Fig. 1G and 1H). In contrast, the inactive YdeH*^GGAAF^*was unable to restore either the phenotype or c-di-GMP level (Fig. 1G and 1H). These results indicated that the c-di-GMP level is directly responsible for the defects in cell size and cell viability observed in the *cdG^0^* strain and that higher c-di-GMP levels did not result in detectable phenotype, consistent with the results obtained in the *cdG^max^* mutant (Fig.1A and 1B). Collectively, these findings indicate that c-di-GMP is essential for maintaining cell size and sustaining cell growth in *Anabaena*.

### Alr2306 is the major PDE to maintain intracellular c-di-GMP for cell growth

During the process of constructing a DGC-null mutant, 12 intermediate mutants with different numbers of genes deleted (1*ΔDGC*, 2*ΔDGC*, 3*ΔDGC*, 4*ΔDGC*, 5*ΔDGC*, 7*ΔDGC*, 8*ΔDGC*, 9*ΔDGC*, 10*ΔDGC*, 11*ΔDGC*, 12*ΔDGC*, 13*ΔDGC*) were generated. Microscopic observations revealed that, in contrast to the other 11 mutants, which exhibited a significant reduction in both cell length and cell width, the 13*ΔDGC* mutant recovered cell length and width similar to the WT strain (Fig. 2A and Fig.S5). Quantitative analysis further demonstrated that the intracellular c-di-GMP level in the 13*ΔDGC* strain was restored to 88 ± 3.7 % of the WT level, whereas the other mutants exhibited c-di-GMP levels ranging from 53 ± 0.79% to 75 ± 6.2% of the WT level (Fig. 2B). Since the 13^th^ gene deleted in the 13*ΔDGC* strain was *alr2306*, and the deletion of *alr2306* alone (1*ΔPDE*) caused a slight increase in c-di-GMP levels (Fig. 1A), we proposed that Alr2306, which contains a receiver (REC) domain at the N-terminal, a GGDEF domain, and an EAL domain at the C-terminal (Fig. S1A), may play a critical role as a major PDE in regulating cell size and growth by modulating c-di-GMP levels.

**Fig. 2.**
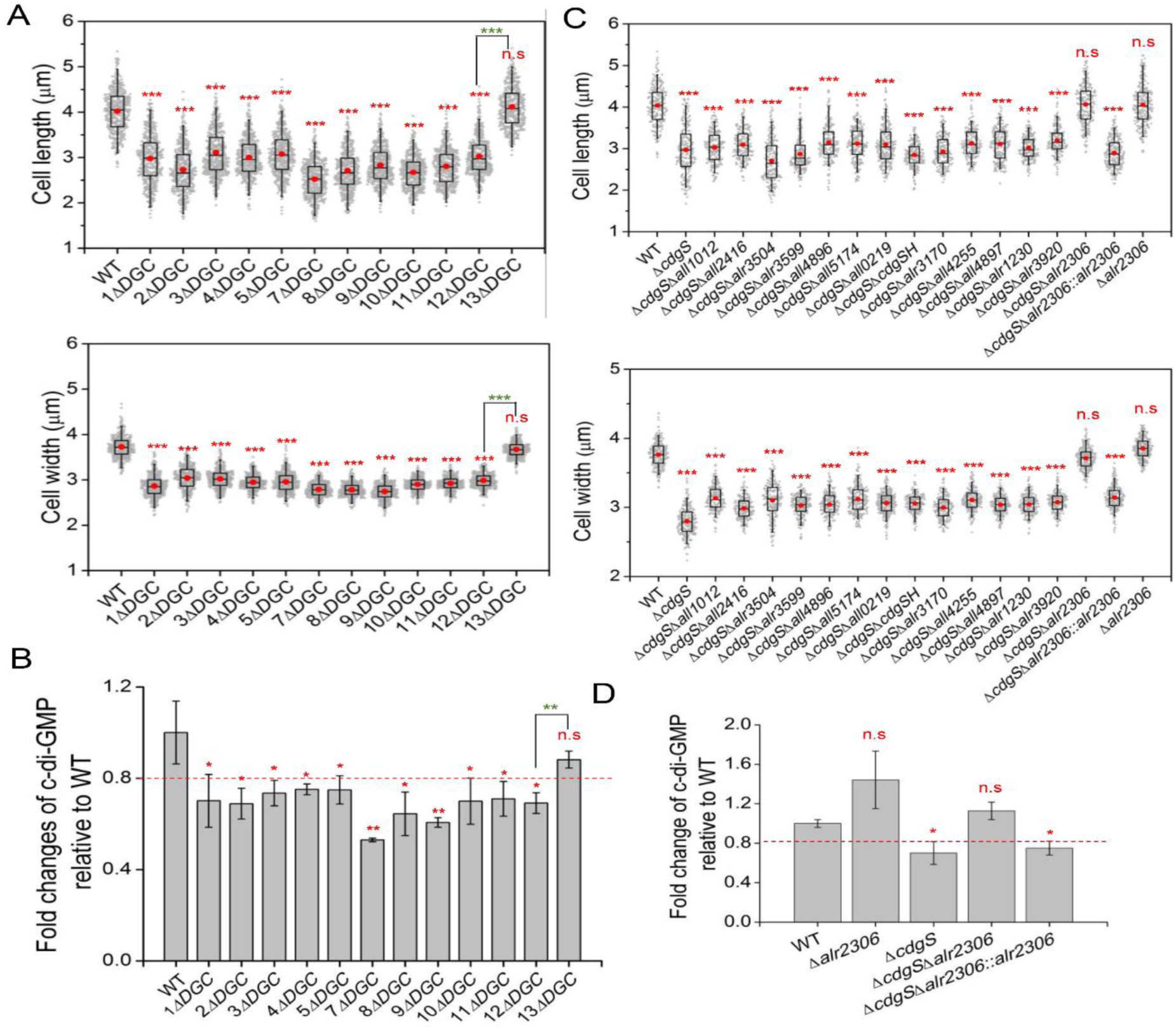
Alr2306 is the major PDE for cell size maintenance. (A and C) Statistical analysis of cell size parameters of the indicated strains. The cell length (upper panel) and cell width (lower panel) were measured based on images, as shown in Fig. S4. 200 cells of each strain from three independent experiments were measured. The boxplots enclose the 25th and the 75th percentile, with the black line representing the median value, red dot representing the mean value. (B and D) The intracellular c-di-GMP level of the indicated strains. The c-di-GMP concentrations of all mutants were normalized to the levels of the wild-type strain. The fold change numbers are shown as mean ± SD from three biological replicates. The red asterisks indicate significance in comparison to WT or between the two strains connected by a bracket at the top as follows: *, p<0.05; **, p<0.01; ***, p<0.001. ****, p<0.000, n.s.: not significant (p >0.05).

To validate this hypothesis, we constructed a series of double mutants by deleting other c-di-GMP metabolic genes in the background of the *ΔcdgS* (1*ΔDGC*) strain, which has been previously identified as the primary DGC involved in cell size regulation (Sun *et al*., 2024). Except for the *ΔcdgSΔall1219* double mutant, we successfully generated 14 other double mutants (Fig.2C and Fig. S5), highlighting the potential importance of *all1219* in c-di-GMP synthesis, which is described later in this study. Among the 14 double mutants, only the deletion of *alr2306* suppressed the reduced cell length and width caused by the deletion of *cdgS* (Fig. 2C and Fig. S5). Consistent with this observation, the intracellular c-di-GMP level of the *ΔcdgSΔall2306* double mutant was also restored to the WT level (Fig.2D). As a control, the complemented strain *ΔcdgSΔalr2306::alr2306* reverted to cell size and c-di-GMP level back to those observed in *ΔcdgS* (Fig. 2C,2D, and Fig. S5). These results demonstrate that the deletion of *alr2306* can elevate the intracellular c-di-GMP levels in *Anabaena* and could partially compensate for the reduced c-di-GMP levels resulting from the loss of *cdgS*. Therefore, although Alr2306 has both a c-di-GMP synthesis domain and a degradation domain, it functions as one of the major PDEs responsible for maintaining intracellular c-di-GMP levels, thereby regulating cell size and growth.

### CdgS, Alr1219, and Alr3599 are the major DGCs that maintain intracellular c-di-GMP levels for cell survival

Because of the initial failure to obtain the *ΔcdgSΔall1219* double mutant, we hypothesized that Alr1219, which contains a CHASE (cyclase and histidine kinase associated sensory extracellular) domain, a transmembrane domain, a PAS (Per-Arnt-Sim) domain, and a GGDEF domain (Fig. S1A), functions together with CdgS as the major DGCs for maintaining cell survival of *Anabaena*. To test this, we generated a conditional strain, *ΔcdgSCT-all1219*, in which the expression of *all1219* is controlled by the CT promoter, as described above. Under induction (0.5 mM TP and 0.3 µM Cu^2+^), the *ΔcdgSCT-all1219* exhibited cell growth and cell size comparable to those of the WT strain, with intracellular c-di-GMP levels approximately 1.2-fold higher than the WT (Fig. 3A-3C). Surprisingly, upon inducer removal, the *ΔcdgSCT-all1219* strain was still able to grow similarly to the WT strain (Fig. 3A and 3E). Microscopic observations revealed that the cell size of the *ΔcdgS CT-all1219* strain initially decreased significantly, but started to recover at 72 h, even back to a cell size similar to the WT strain by 96 h (Fig. 3C). Further quantitative analysis revealed that, after the removal of Cu^2+^ and TP, the intracellular c-di-GMP levels in the *ΔcdgSCT-all1219* strain first decreased dramatically. By 48 h, the intracellular c-di-GMP level dropped to about 42 ± 0.2% of that in the WT strain. However, after 48 h, the intracellular c-di-GMP level began to recover, eventually reaching up to 4-fold higher than that in the WT strain by 96 h (Fig. 3B). The restoration of c-di-GMP levels explains the growth capacity observed in the *ΔcdgSCT-all1219* strain after the inducer removal and suggests that some additional c-di-GMP turnover enzymes may have been activated under conditions when c-di-GMP levels become severely limited.

**Fig. 3.**
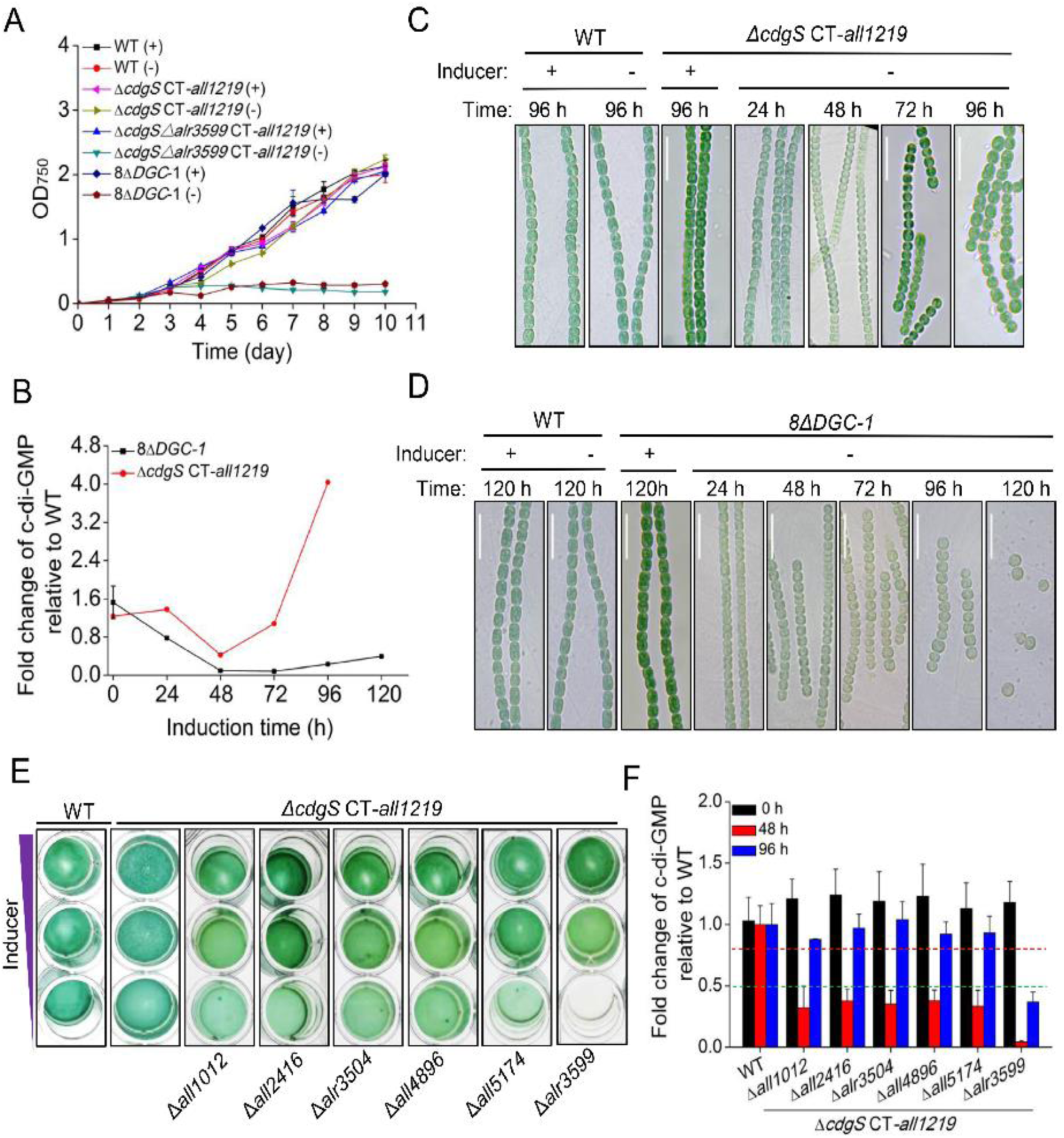
The phenotype and c-di-GMP levels of *cdgS*, *alr3599,* and *alr1219-*related mutant strains. (A) The growth curves of the indicated conditional mutant strains and the WT strain in BG11 medium with (+) or without (-) Cu^2+^ and theophylline. The promoter region of *all1219* in the *ΔcdgSCT-all1219* strain, the Δ8DGC-1 strain, and the *ΔcdgSΔalr3599CT-all1219* strain was replaced by a Cu^2+^ and theophylline inducible platform (CT) at the native chromosomal locus. Absorbance at 750 nm was measured at the indicated time points. All values are shown as mean ± standard deviation, calculated from triplicate data. (B) The intracellular c-di-GMP levels of the *ΔcdgSCT-all1219* strain and the *Δ8DGC-1* strain during the time course of Cu^2+^ and theophylline depletion. (C-D) Micrographs of *Anabaena* filaments of WT, *ΔcdgSpICT-all1219* strains (C), and Δ8DGC-1 strain (D) cultured in BG11 medium with (+) or without (-) Cu^2+^ and theophylline at the indicated time points. Scale bars represent 15µm. (E) The growth of indicated strains was tested in 24-well plates. All cultures started with a similar OD at 0.3 diluted from a pre-culture and imaged after 4 days of incubation. The purple triangle indicated the decreasing levels of the inducers added to the culture media (from up to bottom column): 0.3 μM copper and 2 mM theophylline, 0.125 μM copper and 0.5 mM theophylline, 0 μM copper and 0 mM theophylline. (F) The intracellular c-di-GMP levels of the indicated strains at 0h, 48h, and 96h of Cu^2+^ and theophylline depletion. The green and red dotted line represents the 50% and 80% of the c-di-GMP level to WT, respectively.

To identify these enzymes responsible, we constructed a new mutant *Δ8DGC-1,* in which all 8 genes encoding DGC enzymes alone were deleted (*ΔcdgSΔalr3504Δalr3599 Δall2416Δall1012Δall5174Δall4896CT-all1219*), while the expression of *all1219* is controlled by the CT promoter. Upon removal of the inducer, this *Δ8DGC-1* strain failed to grow, exhibited a significant reduction in cell size, and finally entered growth arrest (Fig. 3A and 3D). These phenotypic observations were supported by intracellular c-di-GMP measurements: following Cu²⁺ and TP depletion, c-di-GMP levels in the *Δ8DGC-1* strain decreased rapidly, becoming undetectable by 48 hours (Fig. 3B). Although a partial recovery of c-di-GMP levels was observed after 72 hours, reaching approximately 40 ± 0.7% of WT levels by 120 h, this c-di-GMP level was insufficient to support cell growth (Fig. 3A and 3B). These results strongly suggest that the activated enzymes in the *ΔcdgSCT-all1219* strain responsible for c-di-GMP restoration are indeed among those absent in *Δ8DGC-1*. Furthermore, these results indicate that the intracellular c-di-GMP level at 40% ± 0.7% of WT is below the threshold required to sustain cell growth, highlighting the critical role of DGC-only enzymes in maintaining c-di-GMP homeostasis and supporting cellular proliferation.

Subsequent triple mutant screening revealed that only the *ΔcdgSΔalr3599CT-all1219* strain exhibited complete cell growth arrest upon removal of Cu²⁺ and TP (Fig. 3E). Quantitative analysis of the intracellular c-di-GMP levels revealed distinct patterns among these mutants (Fig. 3F). The Intracellular c-di-GMP levels in the *ΔcdgSΔalr3599CT-all1219* strain plummeted to around 4.6 ± 0.7% of WT levels at 48 h after inducer depletion and only partially recovered to about 37 ± 8.0% of WT levels by 96 h (Fig. 3F). This limited recovery was insufficient to support cell growth, leading to growth arrest (Fig. 3A and 3E). In the other five triple mutants generated, the c-di-GMP levels dropped to approximately 35% ± 2.5% of the WT levels at 48 h after inducer depletion, and then began to recover thereafter, eventually reaching levels close to the WT by 96 h (Fig. 3G), similar to the *ΔcdgSCT-all1219* strain. This recovery suggests that, if *alr3599* is present, these mutants retain the capacity to restore c-di-GMP homeostasis for cell survival in the absence of CdgS and All1219. Taken together, we concluded that CdgS, Alr1219, and Alr3599 are the major DGCs that maintain intracellular c-di-GMP levels for cell survival, accounting for about 60% of the total c-di-GMP production under the tested conditions. Meanwhile, the limited c-di-GMP level recovery observed in the *ΔcdgSΔalr3599CT-all1219* strain suggests that additional PDE-only and Dual-function enzymes contribute to c-di-GMP level maintenance, responsible for the remaining about 40% of c-di-GMP production in the absence of the CdgS, Alr1219, and Alr3599. Our results also suggest that Alr3599 functions as an emergency DGC to recover the c-di-GMP level independently, once c-di-GMP levels drop below the threshold necessary for cell viability.

### The accumulation of Alr3599, lacking auto-inhibition activity, accounts for its ability to restore c-di-GMP levels under emergency

To characterize the three major DGCs (CdgS, Alr1219, and Alr3599), the Full-length CdgS and Alr3599, along with a truncated Alr1219 variant (Alr1219-ΔCT, lacking both the N-terminal CHASE2 domain and transmembrane domain), were purified from E. coli and assessed for their enzymatic activities. HPLC-based activity assays revealed that all three DGCs exhibited GTP-dependent c-di-GMP synthesis activity (Fig. S6A) (Sun *et al*., 2024). Although all three enzymes followed Michaelis-Menten kinetics, they exhibited significantly different kinetic parameters (Fig. S6B). Most notably, Alr3599 displayed an exceptionally high turnover rate (kcat = 4223.8 ± 1081.1 s⁻¹), approximately 5.3- and 9.2-fold higher than CdgS and Alr1219-ΔCT, respectively. Although Alr3599 showed comparatively lower substrate affinity (Km = 84.2 ± 30.1 μM) compared to CdgS (Km = 26.6 ± 3.5μΜ) and Alr1219-ΔCT (Km = 20.3 ± 12.4 μΜ), its superior turnover rate resulted in the highest catalytic efficiency (kcat/Km = 50.2 ± 12.8 μM⁻¹ s⁻¹) (Fig. S6B). In addition, a pronounced divergence in their regulatory properties was also observed. While Alr1219 and CdgS, containing canonical I-sites (RXXD) (Fig.S1A), were strongly inhibited by c-di-GMP (Ki = 3.7 μM and 2.42 μM, respectively), Alr3599 with its variant I-site (KXXD) showed markedly reduced sensitivity (Ki = 16.0 μM) (Fig. S6C) Taken together, these biochemical characteristics suggested the distinct physiological roles of the three DGCs: Alr3599’s high-output, feedback-resistant design is optimized for rapid c-di-GMP production during emergency responses, while CdgS and Alr1219-ΔCT’s high-affinity, tightly regulated activity maintains precise c-di-GMP homeostasis standard growth conditions.

We next investigated the underlying regulation mechanism of Alr3599-induced activity during c-di-GMP depletion. qRT-PCR analysis showed that the transcriptional level of *alr3599* in the *ΔcdgSCT-alr1219* strain showed no significant changes following inducer depletion (Fig. S6D), despite intracellular c-di-GMP levels dropping to ∼42% of WT concentrations at 48 h (Fig. 3B). However, western blot analysis of a Flag-tagged Alr3599 strain (*ΔcdgSCT-alr1219* (*alr3599-Flag*), Fig.S7) revealed a striking post-translational regulation pattern. Alr3599 protein levels were maintained at barely detectable levels during normal conditions (with inducer), but accumulated dramatically after inducer removal, peaking at 48 h (Fig. S6E). This accumulation correlated temporally with c-di-GMP depletion as a trigger, followed by progressive attenuation of protein levels as cellular c-di-GMP concentrations recovered (Fig. 3B). These results suggest that Alr3599 controls the intracellular level of c-di-GMP at a post-transcriptional level in a c-di-GMP-dependent manner.

### Threshold concentrations of c-di-GMP and phenotypic output in the control of cell size and cell survival

The availability of dozens of mutants constructed during this study and the measurement of their intracellular c-di-GMP levels provided an opportunity to quantify the relationship between the c-di-GMP concentrations and the phenotype observed. We plotted the cell length and width of the WT and all mutant strains exhibiting varying c-di-GMP levels (Fig. 4A). For comparative purposes, the c-di-GMP level of the WT was normalized to 1. The data revealed an S-shaped dose-response relationship between c-di-GMP levels and both cell length and width (Fig. 4A). Analysis of cell length, width, and c-di-GMP concentrations across different mutants, including the genetic consequence of different mutants, identified two distinct function thresholds of the c-di-GMP signal (Fig.4A). First, a morphological threshold: when c-di-GMP levels fall below about ∼80 ± 5% of that of the WT, the cell length and cell width were significantly reduced (Fig.4A). Interestingly, in the range between 80 ± 5% - 50 ± 5%, the output reflected in the reduction of both cell length and cell width is proportional to the drop of c-di-GMP levels, suggesting that c-di-GMP is an important factor for cell size control in *Anabaena*. Second, a viability threshold: A reduction of 50% ± 5% in c-di-GMP levels relative to the WT resulted in complete loss of cell viability (Fig. 4A). These findings establish c-di-GMP as a critical regulator of cell size and survival of *Anabaena*.

**Fig. 4.**
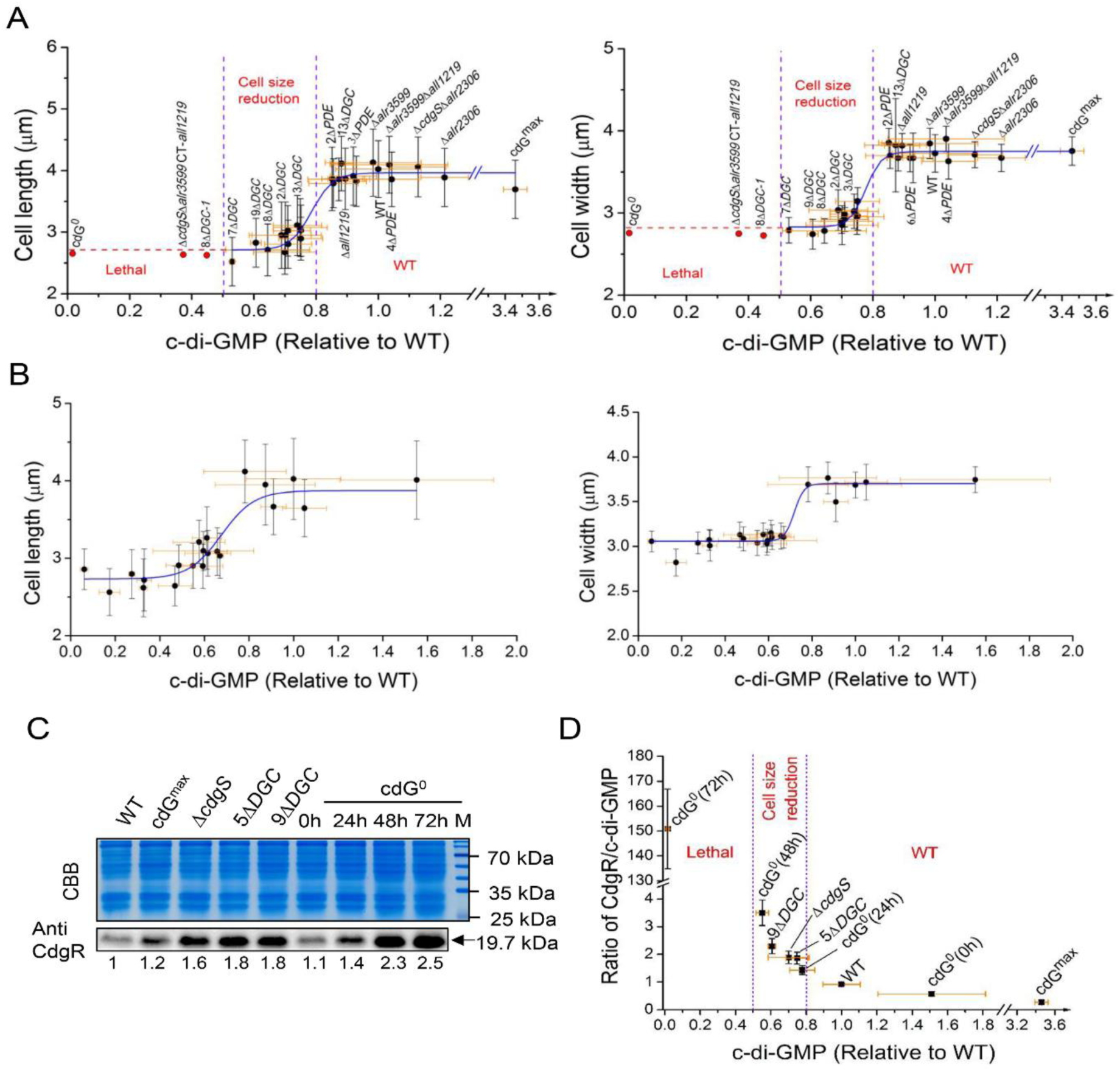
The Interplay among c-di-GMP, CdgR, and cellular morphology in *Anabaena*. (A) Correlation between intracellular c-di-GMP levels and cell length (left) and cell width (right) across different mutant strains. The red dot and dashed line indicate the lethal phenotype of the corresponding strains. Two concentration-dependent regulatory thresholds are indicated by purple dashed lines. (B) Dose-dependent effects of intracellular c-di-GMP levels on cell parameters (cell length (left) and width (right)) in the *cdG^0^* strain under varying concentrations of inducers (Cu^2+^ and theophylline). (C) CdgR abundance in the indicated variant strains and the cdG^0^ strain during the time course of Cu^2+^ and theophylline depletion. Total proteins were visualized with Coomassie Brilliant Blue (CBB), and CdgR was probed with a polyclonal antibody against CdgR (Anti-CdgR). CdgR levels corresponding to those of the WT were quantified with ImageJ from three biological replicates. (D) Correlation between the molar ratios of CdgR to c-di-GMP calculated for the indicated strains and intracellular c-di-GMP levels. In (A) (B) (D), data points represent the mean ± SD from biological replicates.

To further confirm the relationship between c-di-GMP levels and phenotypic outputs, we titrated c-di-GMP concentrations using the conditional cdG^0^ strain and then examined the genetic consequences. As shown in Fig. 4B, the result confirmed the S-shaped dose-response relationship between c-di-GMP concentrations and both cell length and width (Fig. 4B), replicating the two functional thresholds observed in different mutant strains (Fig. 4A). Notably, while the morphological (∼80 ± 5% of WT) and viability (∼50 ± 5% of WT) thresholds remained consistent, this conditional system revealed an additional phenotype: when c-di-GMP levels decreased below 50± 5% of WT, cells first exhibited extreme cell size reduction before progressing to complete cell death, revealing a distinct lag phase progressing to lethal effect, which was not apparent in other mutants. These results provided new insights into the c-di-GMP regulatory functions.

### Cell fate determination by c-di-GMP is dependent on its ability to titrate CdgR

CdgR is a known c-di-GMP-specific receptor that regulates cell size in *Anabaena* (7). Several point mutations in CdgR that abolish c-di-GMP binding lead to cell death (7), suggesting that the increasing abundance of the apo-form CdgR may account for the observed phenotype. To clarify the genetic basis of c-di–GMP–dependent regulation, we deleted *cdgR* in the *cdG^0^* background. This mutation restored both cell size and cell viability to the WT levels (Fig. 1G and 1H), indicating that CdgR mediates the effects of c-di-GMP on these traits in *Anabaena*. We next quantified the intracellular CdgR levels in the WT and mutants with varying cell sizes and c-di-GMP concentrations (Fig. 4C, Fig. S8). In the WT strain, CdgR and c-di-GMP concentrations were 4.44 ± 0.19 μmol/L and 4.84 ± 0.51 μmol/L, respectively, yielding a CdgR/c-di-GMP ratio of 0.92 ± 0.04 (Fig. S8). Since a CdgR dimer binds two molecules of c-di-GMP (7), little apo form CdgR is thus available in the WT strain under normal conditions. In the *cdG^0^* strain, removal of inducers led to a decline in c-di-GMP (see Fig. 1 for details) and a concomitant accumulation of CdgR (Fig. 4C), increasing the CdgR/c-di-GMP ratio (Fig. 4D). When c-di-GMP decreased to 55 ± 11% of the WT, close to the viability threshold, at 48 h after the removal of inducers, the CdgR/c-di-GMP ratio of CdgR/c-di-GMP increased near to 3.5 ± 0.46 (Fig. 4D). By 72 h, c-di-GMP is nearly depleted, with CdgR existing almost entirely in the apo-form (Fig. 4D). Similar results were observed in selected mutants that contain different levels of c-di-GMP and a correspondingly reduced cell size (Fig. 4C and 4D). By cross-examining all the available data (Fig. 1, Fig. 4), we found that the ratio of CdgR/c-di-GMP is about 1.43 ± 0.16 when the c-di-GMP level is at the morphological threshold, and it is about 3.5 ± 0.46 when the c-di-GMP level reaches the viability threshold. Thus, the ratios of CdgR to c-di-GMP act as a central determinant of the phenotypic output through a titration mechanism. These results support our previous hypothesis that the accumulation of the CdgR apo form ultimately determines cell fate in *Anabaena* (7).

## Discussion

Most bacterial genomes encode numerous enzymes for c-di-GMP synthesis and degradation (Agostoni *et al*., 2013; Chen *et al*., 2020; Solano *et al*., 2009). However, the mechanisms governing c-di-GMP pool dynamics remain poorly characterized, particularly in cyanobacteria. *Anabaena* possesses 16 c-di-GMP metabolic enzymes and has been reported to utilize c-di-GMP as a central regulator for cell size control (Huang *et al*., 2021; Sun *et al*., 2024). In this study, we created a large number of mutants, including two (*cdG^max^*and *cdG^0^*) in which all genes encoding c-di-GMP synthesis and degradation enzymes were deleted, respectively (Fig. 1 and Fig. S1-S4). Due to the relatively slow growth rate of cyanobacteria such as *Anabaena*, the deletion of such a large number of genes in one single mutant was unreported in cyanobacterial genetics. This long-term genetic effort led us to make several important conclusions on the function of the c-di-GMP signal in *Anabaena*. Firstly, our data indicate that c-di-GMP serves not only as a modulator for cell size control but also as an essential signaling molecule required for cell viability in *Anabaena* (Fig.1 and Fig.S1-S4). Secondly, quantitative analysis revealed two physiologically relevant c-di-GMP thresholds in vivo: a minimal level necessary for cell size maintenance and a critical lower limit required for cell viability (Fig. 4). Thirdly, the 16 c-di-GMP turnover enzymes in *Anabaena* can be classified into different categories based on their contribution to the pool of c-di-GMP (Fig. 5). Among all 16 c-di-GMP metabolic enzymes, the previously identified CdgS (Sun *et al*., 2024), All1219, and Alr3599 act as the primary DGCs (Fig.3 and Fig.S6), while Alr2306 serves as the primary PDE (Fig. 2). These enzymes form a core enzymatic network that orchestrates c-di-GMP homeostasis to regulate cell size and viability (Fig. 5). It also revealed a safeguard mechanism with Alr3599, which is induced post-transcriptionally to rescue the level of c-di-GMP when it drops below the viability threshold (Fig. 3). Our results also demonstrate that both the cell size control by, and the essential function of, c-di-GMP are dependent on its ability to saturate CdgR, corresponding to the two threshold concentrations, and thus the two ratios of CdgR/c-di-GMP (Fig. 4). Part of the functions of c-di-GMP-CdgR signaling pathway is mediated by DevH, an essential transcription factor in *Anabaena* (Xu *et al*., 2025; Zeng *et al*., 2023). Both DevH and NtcA are transcription factors belonging to the CRP family. The DevH regulon strongly overlaps that of NtcA and controls a multitude of cell physiology (Xu *et al*., 2025). Other downstream factors, in addition to DevH, are likely involved in the control of cell size and viability.

**Fig. 5.**
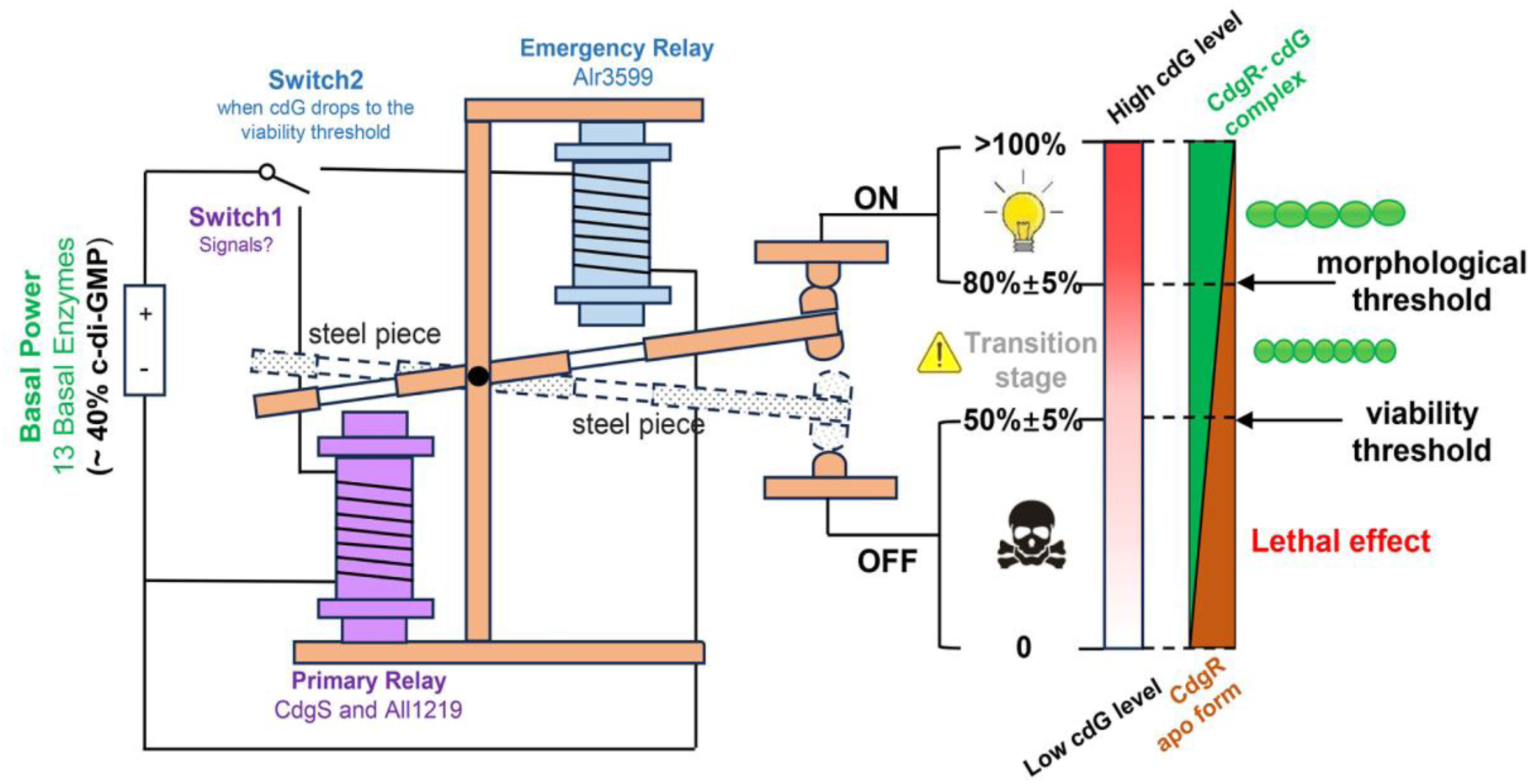
A c-di-GMP dual relay system governs cell growth and survival in *Anabaena*. The system consists of three functional modules: (1) Basal power supply: Maintains a basal c-di-GMP level (∼ 40%) through the activity of 13 enzymes, including the dominant PDE Alr2306. This module is analogous to a battery that powers both relays and provides a baseline signal amplification. (2) Primary Relay Unit (Switch1): Comprises the DGCs CdgS and All1219, which operate like an electromagnetic coil. Their contribution (∼60% of total c-di-GMP under standard conditions), which depends on the input signals, combined with the basal c-di-GMP, determining three output states: “ON” state (>80 ± 5% c-di-GMP): Maintains normal *Anabaena* morphology (steel piece engaged); Transition state (50 ± 5 - 80 ± 5% c-di-GMP): Induces cell size reduction proportional to c-di-GMP level; “OFF” state (<50% ± 5 c-di-GMP): Cause lethal effect of *Anabaena* (steel piece disengaged). (3) Emergency Relay Unit (Switch 2): The dormant DGC Alr3599 activates when c-di-GMP drops to the viability threshold (50 ± 5% c-di-GMP), functioning as a backup coil, it triggers steel-piece rearrangements to force the “OFF” state to return to the “ON” state, saving cells from death. A dual-threshold control mechanism (morphological threshold: 80±5%; viability threshold: 50 ± 5%) ensures precise regulation of cellular c-di-GMP homeostasis. The phenotypic outcomes, including reduced cell size and lethality, are ultimately determined by the molar ratio of CdgR to c-di-GMP. Arrows indicate threshold values. cdG: c-di-GMP

Based on these findings, we propose an innovative regulatory model in *Anabaena,* where the 16 c-di-GMP metabolic enzymes integrate into a sophisticated and responsive dual-threshold control system for c-di-GMP level in vivo (Fig. 5). This system dynamically maintains c-di-GMP homeostasis to coordinate cell size and viability through two concentration thresholds, with cell size reduction and loss of cell viability as outputs. This biological control system operates on principles analogous to electromechanical dual-relay circuits, but with genetic consequences as biological outputs (Fig. 5). Based on their contribution to the c-di-GMP pools, the 16 c-di-GMP metabolic enzymes can be classified into three categories, corresponding to key electromechanical relay modules: basal power, primary relay, and emergency relay (Fig. 5).

The enzymes of category 1 constitute the basal power module. This module contributes to the foundational and constitutive c-di-GMP pool (∼40%), which is maintained by 13 enzymes, including the dominant PDE Alr2306. This module is analogous to a battery that powers both relays and provides a baseline signal amplification regardless of which switch dominates (Fig. 5). Enzymes of category 2 constitute the primary relay unit (Switch 1), and they include the most important DGCs CdgS and All1219. This module functions as an electromagnetic coil in the primary relay, contributing to around 60% of cellular c-di-GMP under standard tested conditions (Figs. 5 and 3B). CdgS and CdgK constitute a two-component system, and CdgK possesses multiple sensor domains at the N-terminal (Sun *et al*., 2024). Alr1219 also has CHASE and PAS sensor domains (Fig. S1A). These observations suggest that the contribution of CdgS and Alr1219 could depend on input signals. When total c-di-GMP (Basal power + Primary Relay) exceeds the morphological threshold (80±5%), this module signals an “ON” state through a system that simulates the iron-core-mediated steel-piece conformational changes, producing standard cell morphology. When total c-di-GMP is at intermediate levels (50 ± 5 – 80 ± 5%), this module is in a transition state, producing reduced cell size proportional to c-di-GMP level. When total c-di-GMP is under the viability threshold (50±5%), this module is in an “OFF” state, leading to a lethal effect. The third category acts as an emergency relay module (Switch 2). This module centers on the DGC Alr3599, a dormant sensor that activates only when c-di-GMP plummets to below the viability threshold (50±5%). Like a backup coil, it triggers a cell response in a manner comparable to a steel-piece rearrangement to force transition from an “OFF” to “ON” state, thus preventing cell death. The different c-di-GMP threshold concentrations translate into corresponding c-di-GMP/CgdR ratios, which ultimately dictate cell fates and responses (Figs. 4 and 5). Certain details in this model system still need to be understood in particularly the specific signals perceived by the CdgK-CdgS and Alr1219 sensor domains that modulate cellular c-di-GMP levels and their regulation mechanism, and the precise post-transcriptional activation mechanism of Alr3599 under c-di-GMP-limited conditions. Nevertheless, this biological dual relay system, integrating basal maintenance, signal-responsive modulation, and emergency override, achieves both precise and robust control of cellular c-di-GMP homeostasis. Employing threshold-dependent state transitions resolves the fundamental growth-survival trade-off that challenges most living systems.

In bacteria, multiple signaling pathways can operate concurrently to generate distinct physiological outputs while utilizing the same diffusible second messenger maintained at a consistent global concentration (Hengge, 2021; Junkermeier & Hengge, 2023). This remarkable signaling specificity arises from the integration of local and global c-di-GMP signaling modalities within complex regulatory networks (Junkermeier & Hengge, 2023). Our study reveals that c-di-GMP regulates the cell size and viability in *Anabaena* (Fig. 1). Unlike well-characterized systems where c-di-GMP mediates different cellular processes through local c-di-GMP signaling (Junkermeier & Hengge, 2023; Junkermeier & Henggea, 2021; Richter *et al*, 2020; Sellner *et al*, 2021; Thongsomboon *et al*, 2018) (e.g., the c-di-GMP-dependent Bcs exopolysaccharide-producing system (Richter *et al*., 2020; Thongsomboon *et al*., 2018) and the Nfr vicinity system in E. coli (Junkermeier & Henggea, 2021; Sellner *et al*., 2021)), our data suggest that *Anabaena* primarily employs global c-di-GMP signaling to coordinate the cell size and viability (Fig. 1C and 1H). Several lines of evidence support this conclusion. First, while targeted deletion of the diguanylate cyclase CdgS produces a specific phenotype, the accompanying substantial changes in total cellular c-di-GMP levels strongly implicate a global signaling mechanism (Fig. 1C). Second, both cell size and viability regulation converge on the same c-di-GMP effector, CdgR (Fig. 1H), which shows no physical interaction with pathway-specific DGCs (Fig. S9). Notably, *Anabaena* also exhibits a distinct c-di-GMP regulatory logic compared to canonical paradigms. In most bacteria, elevated c-di-GMP levels promote sessile behaviors (Li *et al*, 2022) (Yam *et al*, 2022) (Chen *et al*, 2023)(e.g., biofilm formation in *Pseudomonas aeruginosa* (Katharios-Lanwermeyer *et al*, 2021)). However, in *Anabaena*, elevated c-di-GMP levels did not induce phenotypic changes at test conditions, whereas c-di-GMP reduction triggered an adaptive response in cell size control (Fig. 1) (Sun *et al*., 2024). This inverse correlation suggests a fundamentally different wiring of c-di-GMP signaling networks in this organism.

Cell size homeostasis represents a fundamental biological characteristic across bacterial species (Chien *et al*, 2012; Taheri-Araghi *et al*, 2017). Recent work has established ppGpp as a central regulator of bacterial cell size, directly linking metabolic status to division control through growth rate modulation and adder-based mechanisms in *E. coli* (Büke *et al*, 2022). Strikingly, our studies reveal that *Anabaena* employs c-di-GMP, not ppGpp, as the primary determinant of cell size, while retaining functional ppGpp synthetases/hydrolases (Fig. S10). Overexpressing or disrupting the single relA/spoT homolog protein All1549 in *Anabaena* (strain OE-*all1549* or *all1549Ωsp/sm*) failed to alter cell size significantly (Fig. S10)(Zhang *et al*, 2013), while perturbations in c-di-GMP pools elicited pronounced cell size effects (Fig. 1). The observed divergence in cell size regulatory systems likely stems from unique physiological constraints imposed by the photosynthetic lifestyle and multicellular complexity of *Anabaena*. Except partly the roles of the DevH (Xu *et al*., 2025; Zeng *et al*., 2023), the molecular mechanisms by which these second messengers govern cell size remain to be fully elucidated and represent an important direction for future investigation.

In summary, this study demonstrates how nature repurposes the conserved signaling molecule c-di-GMP to construct novel regulatory architectures adapted to specific environmental contexts. Beyond revealing novel adaptation strategies in cyanobacteria, this mechanism provides a blueprint for engineering synthetic biological systems that require multi-level regulation.

## Methods

### Bacterial strains, media, and growth conditions

*Anabaena* sp. PCC 7120 and its derivatives were cultivated at 30°C in BG11 medium under continuous illumination (30 μmol m^−2^.s^−1^) with shaking at 180 rpm (Xing *et al*, 2022). For conditional mutants, where gene expression was controlled by the artificial CT promoter (Xing *et al*, 2021), cells were initially grown in BG11 medium supplemented with 0.3 μM copper and 0.5∼1.0 mM theophylline (TP) to induce gene expression during the logarithmic phase. To terminate gene expression, cells were washed three times with a BG11 medium devoid of copper and theophylline and then transferred to a fresh copper/TP-free BG11 medium. When required, Neomycin (25 μg/ml), streptomycin (2.5 μg/ml), or spectinomycin (5 μg/ml) was provided in the culture. All strains used in this study are listed in Table S1.

### Construction of plasmids and cyanobacterial recombinant strains

All markerless deletion mutants, conditional mutants, and strains with point mutation were created using the genome editing technique based on CRISPR-Cpf1 as previously described (Niu *et al*., 2019; Xing *et al*., 2021). Plasmids containing the artificial CT promoter for inducible gene expression (using copper and theophylline) were constructed based on the PCT vector. For protein expression in *E. coli*, the corresponding DNA fragments were amplified and sub-cloned into the pHTwinStrep vector containing two carboxyl-terminal Strep-tags at the C-terminal. Plasmids containing genes with point mutations were generated using quick-change mutagenesis, with the plasmid harboring the corresponding wild-type (WT) gene as the template.

Markerless deletion and overexpression strains were created by conjugating the corresponding plasmids into *Anabaena (Zeng et al., 2023)*. For the *all1219* conditional mutant, the native promoter region of *all1219* was replaced with the artificial CT promoter. Point mutation strains were constructed as previously described (Sun *et al*., 2024). All primers and plasmids used in this work are listed in Table S2 and Table S3, respectively.

### Estimation of Intracellular Concentration of CdgR and c-di-GMP

The packed cell volume was determined as previously described with minor modifications. Briefly, 9 mL of WT cell suspension (OD_750_=0.462) was added to a hematocrit tube, and the sample was centrifuged in a hematocrit tube at 5,000 × g for 30 minutes, yielding a packed cell volume of 10.75 μL. This value corresponds to a normalized packed cell volume of 2.58 μL/OD₇₅₀, which is consistent with results reported by Laurent et al. The cell volume was also estimated geometrically by modeling the cell as a cylinder, using the formula: V= π × (1/2 cell width) × (1/2 cell width) × cell length. To quantify the cellular abundance of CdgR, an immunoblot assay was employed. Cell extracts normalized to 0.3 OD₇₅₀ units were resolved by SDS-PAGE alongside a standard curve generated from a dilution series of purified CdgR protein of known concentration. Following transfer, the membrane was probed with specific anti-CdgR antibodies. The band intensities from the cell lysates were interpolated against the standard curve to determine the absolute amount of CdgR. The concentration of CdgR and c-di-GMP was then calculated based on their quantification and the corresponding cell volume and cell numbers, with the final value representing the mean of three independent biological replicates.

### Protein expression and purification, extraction and quantification of cellular c-di-GMP, in vitro assays of DGC activity, quantitative real-time PCR (qRT-PCR), and western blots

Details of protein expression and purification, extraction and quantification of cellular c-di-GMP, in vitro assays of DGC activity, quantitative real-time PCR (qRT-PCR), and western blots were reported previously (Huang *et al*., 2021; Sun *et al*., 2024; Zeng *et al*., 2023). Detailed descriptions are provided in the SI Appendix.

### Image acquisition and cell size analysis

The microscope images were captured by the SDPTOP EX30 microscope and processed using ImageJ without deconvolution as described previously (REF). For cell size analysis, microscopy images were taken when the optical density of a culture reached 0.3-0.5 at OD_750_. The cell length and width of cells of each strain were measured by ImageJ. Statistical tests and plotting of data were performed with Origin 8.0.

### Statistics and reproducibility

All statistical analyses and data visualization were performed using SPSS software version 19.0 (SPSS Inc.) and Origin software, respectively. Each data point in the plots represents an independent measurement derived from at least two or three biologically independent experiments. Comparison between the control and tested groups was performed using an independent sample t-test. Multiple comparisons were done using One-Way ANOVA followed by Sidak’s multiple comparisons test. Significance thresholds (p-values) are provided in the figure legends.

## Acknowledgments

We would like to thank Jun Men and Siyu Wang from the Analysis and Testing Center of the Institute of Hydrobiology, Chinese Academy of Sciences, for support in the measurement using HPLC and LC–MS/MS. This research was supported by the Youth Innovation Promotion Association CAS (No. 2022342), the National Natural Science Foundation of China (Grant No. 32270063), the Strategic Priority Research Program of the Chinese Academy of Sciences (Grant No. XDB0480000), and the China Postdoctoral Science Foundation (Grant No. 2023M743726).

## Author contributions

Conceptualization, CZ.Z., and X.Z.; Methodology, QX.S.; Investigation, QX.S. and M.H.; Formal Analysis, QX.S., and X.Z.; Visualization, QX.S. and X.Z.; Writing – Original Draft: X.Z., Writing – Review & Editing: X.Z. and CC.Z.; Funding Acquisition: CC.Z., X.Z., and QX.S..

## Data and materials availability

Data are available in the published article and its online supplemental material. Plasmid pCpf1b-sp was deposited at Addgene with ID number #122188. Plasmid pCT was submitted to GenBank with ID number MK948095. For any further inquiries about the work, please contact the corresponding author.

